# Dark exposure reduces high-frequency hearing loss in C57BL/6J mice

**DOI:** 10.1101/2024.05.02.592252

**Authors:** Peter Jendrichovsky, Hey-Kyoung Lee, Patrick O. Kanold

## Abstract

Plastic changes in the brain are primarily limited to early postnatal periods. Recovery of adult brain plasticity is critical for the effective development of therapies. A brief (1-2 week) duration of visual deprivation (dark exposure, DE) in adult mice can trigger functional plasticity of thalamocortical and intracortical circuits in the primary auditory cortex suggesting improved sound processing. We tested if DE enhances the ability of adult mice to detect sounds. We trained and continuously evaluated the behavioral performance of mice in control and DE conditions using automated home-cage training. Consistent with age-related peripheral hearing loss present in C57BL/6J mice, we observed decreased performance for high-frequency sounds with age, which was reduced by DE. In CBA mice with preserved peripheral hearing, we also found that DE enhanced auditory performance in low and mid frequencies over time compared to the control.

## Introduction

Plastic changes in the brain are primarily limited to early postnatal periods and have been mainly studied within the same modality. Crossmodal plasticity refers to neural plasticity that allows adaptation to the loss of a sensory modality. Sensory modality loss can occur during pathological states of the peripheral sensory systems, such as loss of hair cells in the cochlea, resulting in deafness, or damage of the eyes, resulting in blindness (1-4). Crossmodal plasticity is thought to underlie the enhanced auditory abilities of the early- (1, 5-8) and late-blind (7, 9). Several circuit-level plasticity changes have been observed in adult mice with temporary visual deprivation (dark exposure, DE). DE in adult mice has profound effects on the primary auditory cortex, such as strengthening of thalamocortical synapses (10), refinement of excitatory and inhibitory intracortical circuits (11-13), and selective reduction of thalamic-reticular nucleus-mediated inhibition of the auditory thalamus (14). These circuit changes correlate with changes in the sound-evoked responses, such as reduced thresholds, increased gain, increased frequency selectivity (10), and decorrelation of spatiotemporal population responses (15), which together should lead to increased coding fidelity. We thus investigated if DE leads to improved auditory ability.

To understand the effects of DE on auditory processing, we placed 48 C57BL/6J (C57) and 48 CBA/CaJ (CBA) mice in home cages fitted with an automated auditory behavior system (“ToneBox”) that allowed continuous long-term observation (Fig. 1A) (16, 17). To test the effects of DE, we take advantage of both the C57BL/6J mouse strain that develops progressive hearing loss with age as well as CBA mice that retain normal hearing (18-24). Using the C57BL/6J strain enables us to test the effects of DE on hearing frequency bands with normal (low and mid frequencies) and reduced complement of hair cells (high-frequencies).

**Figure 1:**
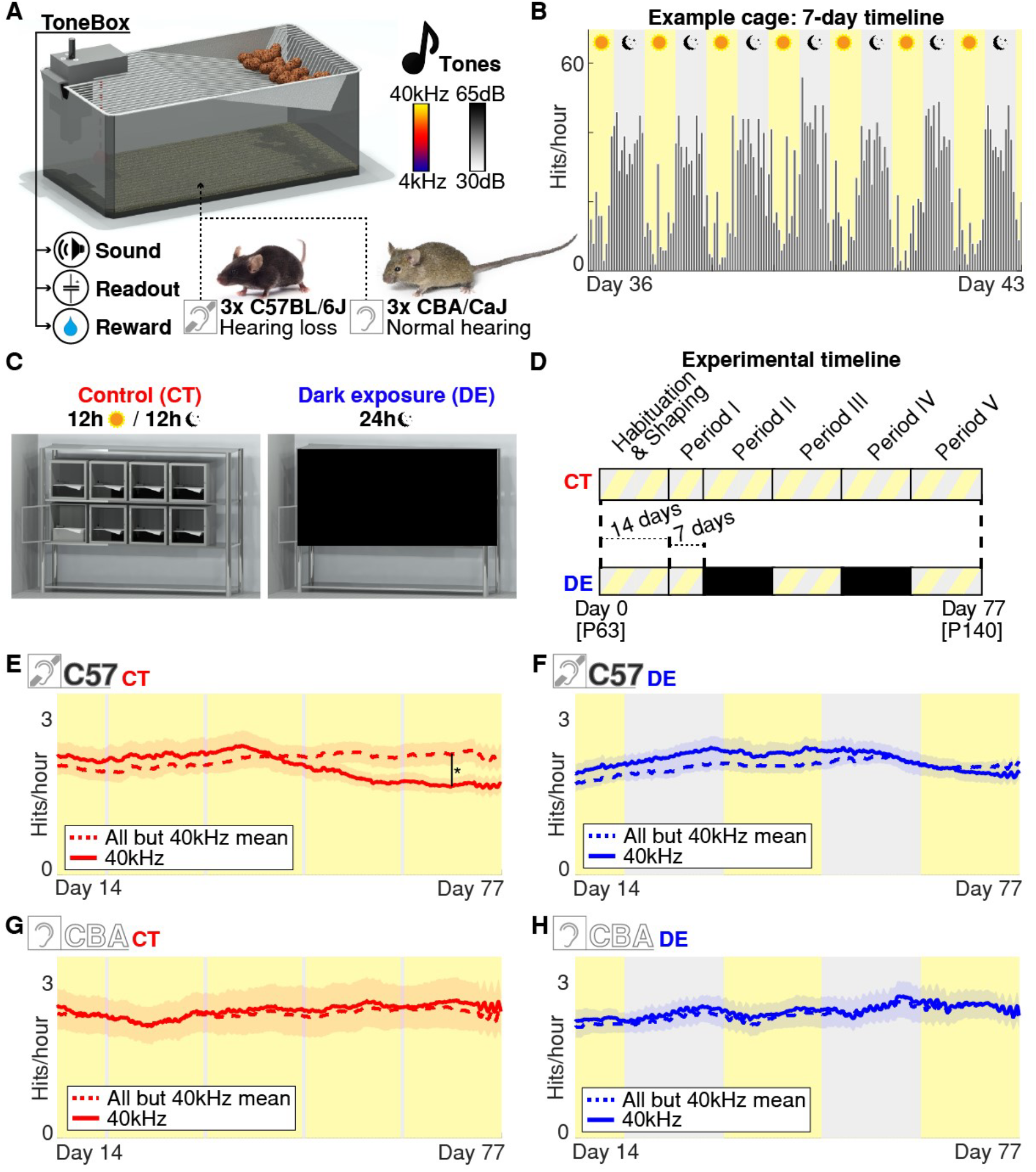
DE reduces loss of detection performance for high-frequency tones. **(A)** Automated home-cage training system with ToneBox. Tones are randomly presented from 4-40kHz and 30-65dB amplitude. Three animals were placed in each training cage, either C57BL/6J or CBA mice. These groups are labeled with a crossed-out ear pictogram for the C57BL/6J group and a non-crossed-out ear pictogram to label the CBA group. This notation is used for all figures. **(B)** Example performance during one week. Gray bars indicate the hit rate within each hourly time bin. **(C)** 8 ToneBoxes under control 12h/12h light/dark conditions or during DE. **(D)** Experimental timeline. DE cohort receives two 2-week DE periods (II & IV). Animals begin the Habituation and Shaping phase at postnatal day 63 (P63). **(E, F)** Moving average hit rates for 40kHz or all other tones for C57BL/6J CT and DE groups (N=8 for both). The black line in (E) on day 70 shows the difference between the two observed means and this difference is statistically significant (*t-test, p* < 0.05). Data averaged over all sound amplitude levels. **(G, H)** Same as in (E, F) but for CBA CT and DE groups (N=8 for both).

Once placed in the ToneBox, mice received a water reward for detecting tones from the ToneBox speaker. After the tone was presented, mice had a reward window to lick the ToneBox water spout. Licking was detected with a capacitive sensor. By placing multiple animals in the cage, we avoided the effects of social isolation. Mice lived and performed the task in the cage with no interaction with humans for the duration of the experiment, except for biweekly bedding change. In the ToneBox, we continuously presented 88 different combinations of different sound frequencies (11 total tones) and sound amplitudes (8 total amplitudes), which enabled us to construct long-term “performance audiograms.” Figure 1B shows an example of a typical week-long timeline of hit activity showing, as expected, that mice were active in the dark cycle.

## Results

### DE reduces the decline in performance in the high-frequency band of C57BL/6J mice

Mice were 63 days old at the start of the experiment. We divided the mice of both C57BL/6J and CBA lines into two groups respectively: Control (CT) and DE (8 ToneBoxes, 24 mice per group) (Fig. 1C). Mice were placed into the ToneBoxes for an initial 14-day habituation and shaping phase to stabilize their performance (Fig. 1D). We divided the following 70-day-long experiment timeline into five periods. Period I was 7 day-long while Periods II, III, IV, and V were 14 day-long. Periods I, III, and V were under a typical 12-hour light/dark cycle for both experimental groups. In periods II and IV, the DE group was hermetically sealed from the room light, while the CT group remained in the normal Light/Dark cycle. Normal circadian rhythm was present throughout the experiment, including DE periods. Example recordings of hit rates from the complete 77-day-long timeline are shown for all four groups: the C57BL/6J CT, CBA CT, C57BL/6J DE, and CBA DE groups (Fig. S1).

In CT C57BL/6J ToneBoxes, we noticed that the hourly hit rates show a decrease in performance for the highest frequency band (40kHz) with increasing age (Fig. 1E). We performed a spot test on day 70 of the 40kHz band against performance from all other frequency bands and the difference was statistically significant (*t-test, p* = 0.0273). This decreased performance is consistent with the development of presbycusis in these mice due to peripheral hearing loss (18-20). Based on ABR measurements (18-20), the onset of hearing loss in C57BL/6J mice in the high frequencies is approximately at the start of Period II (Postnatal day 84, P84), consistent with the decreased performance we observed. In contrast, DE C57BL/6J ToneBoxes did not show decreased performance at high frequencies and only showed a slight decline later in the experimental timeline (Fig. 1F). This delay in decline will be described below in detail. At day 70 performance was similar between groups (*t-test, p* >> 0.05). Consistent with preserved peripheral hearing in CBA mice, the 40kHz band did not show any decline with age, and groups performed similarly at day 70 (*t-test, p* >> 0.05) (Fig. 1G and 1H). These observations suggest that the DE periods in visually deprived animals reduced the decline of the processing of high-frequency sounds in C57BL/6J mice.

### DE increases tone detection performance in CBA and C57BL/6J mice

To first identify how animal performance varied with frequency and amplitude and to investigate whether DE affected the frequency and amplitude (FxA)-dependent hit rate, we calculated the ratio of hit rates between the first vs. last period hit rates for each FxA bin. We first plotted these heatmaps for C57BL/6J CT and DE groups (Fig. 2A). In the CT C57BL/6J group, the hit rates decreased for the whole high-frequency band across sound amplitudes (Fig. 2A, black arrows, 32 & 40kHz all SPL levels). This decline in performance occurs in a similar frequency band as presbycusis reported in C57BL/6J mice of this age (19). In contrast, such decreased performance was not present in the C57BL/6J DE group. These results suggest that DE attenuates the high-frequency-specific decline of detection performance in C57BL/6J mice.

**Figure 2:**
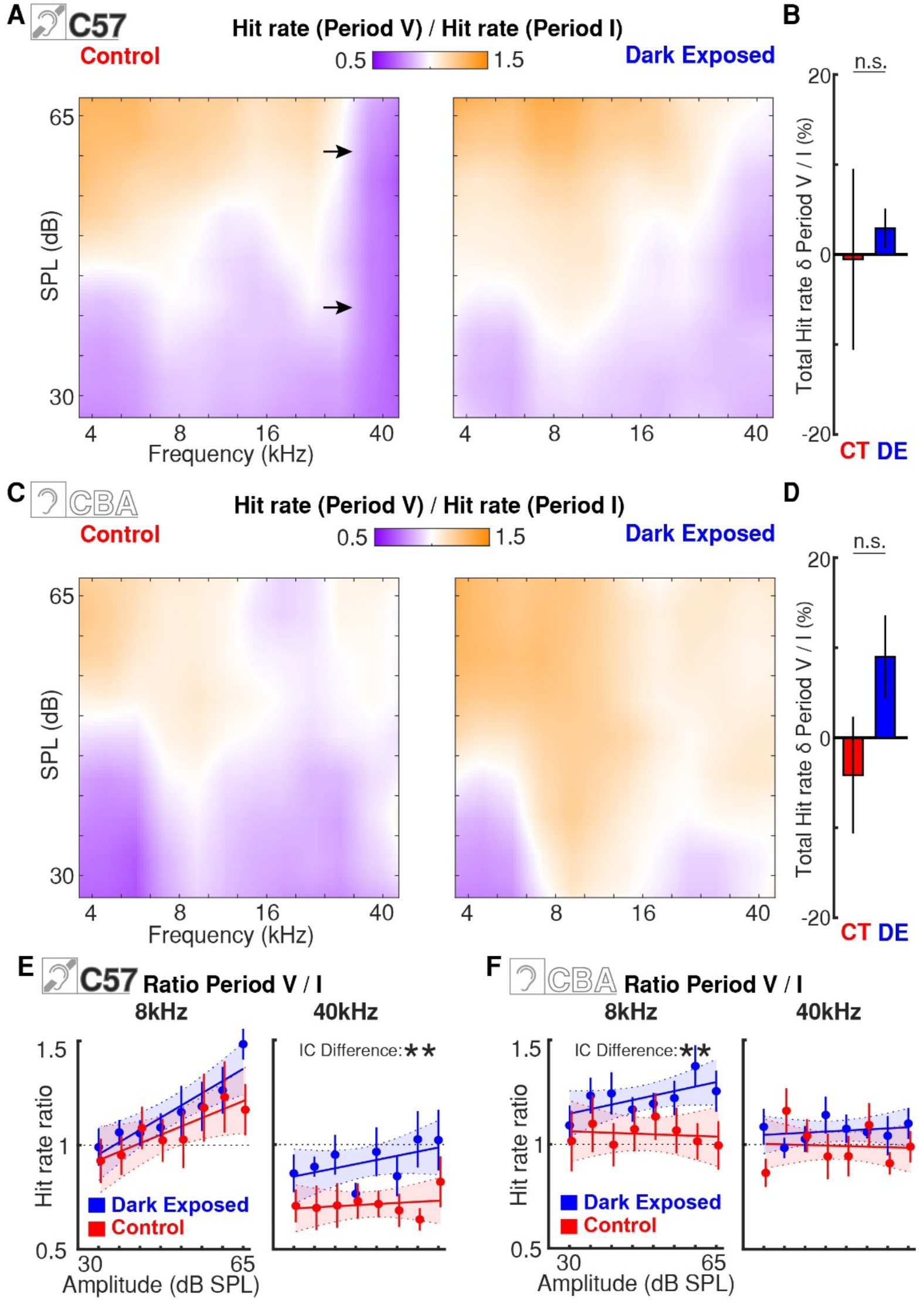
DE causes broad increases in performance in CBA and C57BL/6J mice. **(A)** Rate-of-change audiograms for the CT (Left) and DE (Right) C57BL/6J groups. Black arrows indicate the area of high-frequency band performance which significantly deteriorated throughout the experiment. **(B)** Bar plot showing the average change of total period hit rates between periods I and V. Red C57BL/6J CT, Blue DE C57BL/6J group. Black lines indicate SEM. ‘n.s’, ‘*’, indicate statistical significance (*t-test*, non-significant, *p* < 0.05 respectively). **(C)** Rate-of-change audiograms for the CT (Left) and DE (Right) CBA groups. **(D)** Same as in B, but for CBA CT and CBA DE groups. **(E)** Scatter points show hit rate ratios for 8kHz and 40kHz as a function of amplitude in the C57BL/6J CT (red) and DE (blue) groups between Periods I and V. Vertical lines show SEM. Scatter points are overlapped with linear regression model fit in matching colors. Shaded areas are 95% confidence intervals of the fit. Intercept (IC) or slope (SL) Difference appears if the *p*-value of the *F*-test showed significance for either parameter of the group difference linear fit model (*F*-test, Bejnamini & Hochberg false discovery rate procedure applied. Does not appear for non-significant, * for *p* < 0.05, & ** for *p* < 0.01). Dashed black horizontal lines outline an NHR level of 1. Dashed black horizontal lines outline hit rate ratio 1 (Where mean performance in both periods was the same) **(F)** Same as in (E) but for CBA CT and DE groups.

We next investigated if there was an overall decreased performance in CT C57BL/6J mice compared to DE mice. We calculated the total hit rates by merging all FxA bins in a given period, which gave us a rough measure of change in overall tone detection performance, independent of the tested stimulus (Fig. 2B, left). The changes in total hit rates in CT and DE C57BL/6J mice were similar (CT: -0.53% ± 10.06% SEM, DE: + 2.91% ± 2.19% SEM, *t-test, p* = 0.7428). This indicates that the overall performance of CT and DE mice was similar. Given that hit rates decreased for high frequencies in CT mice, this suggests that CT mice have relatively more hits at lower frequencies. In contrast, in C57BL/6J DE mice high-frequency performance is preserved. Together, these results suggest CT C57BL/6J mice show decreased hit rates for high frequencies and an increased hit rate at lower frequencies while in DE C57BL/6J mice high-frequency performance is preserved.

We next analyzed the rate-of-change audiograms for the CBA CT and DE groups (Fig. 2C). In contrast to C57BL/6J mice CT CBA mice do not show decreased performance at high frequencies. However, after DE we observed a widespread increase in hit rates (Fig. 2C). In the CBA DE group hit rates increased by +9.00% ± 4.60% SEM while in the CBA CT group hit rates decreased by -4.15% ± 6.49% (Fig. 2D, left; *t-test, p* = 0.1205). While this difference is not significant when summed over the whole frequency spectrum, qualitative inspection suggests that the frequency band-specific performance in low and mid frequencies is enhanced by DE.

To examine band-specific performance for both C57BL/6J and CBA groups in detail we analyzed the relationship between sound amplitude and hit rates in each frequency band. All trials for the C57BL/6J CT group were first binned based on the sound amplitude parameter, and the global correlation coefficient was calculated to be 0.9679 ± 0.0058 SEM, while the remaining three groups followed a similar pattern. We thus used linear regression analysis to test whether the slope (gain, abbr. SL) of this relationship and/or baseline performance (intercept, abbr. IC) is changed after DE for C57BL/6J mice (Fig. 2E) and CBA mice (Fig. 2F). We used linear regression model fitting to estimate the parameters of the linear fit for each frequency band and each group. These linear fit estimates, together with 95% confidence intervals (CI) are shown as lines in respective colors (fit) and 95% CI as shaded areas for plots in both subpanels. To determine if these model estimates differ, the difference between the two groups was also fitted and tested for the significance of parameters using a Multiple comparison correction (Bejnamini & Hochberg false discovery rate procedure (25)). In Fig. 2E and 2F, we plot results for 8kHz and 40kHz bands, and the statistics for the remaining frequency bands are given in Table S1. First, this analysis confirms our visual observation of a major decline in the 40kHz band for the C57BL/6J group in Fig. 2A where IC of CT and DE fits differed significantly (*F*-test, *p* = 0.0027). Secondly, the frequency band-specific comparisons for CBA mice show differences of the intercept parameter between the two groups for several low- and mid-frequency bands (4kHz: p = 0.0115; 6.3kHz: p = 0.0014; 8kHz shown in Fig. 3B: p = 0.0079; 10kHz: p = 0.0005; 12.5kHz: p = 0.0107; 16kHz: p = 0.0071; 25.0kHz: p = 0.0107; all other n.s. bands in Table S1). These rate-of-change audiograms suggest that DE increased the total period hit rate across a broad frequency range in CBA mice.

**Figure 3:**
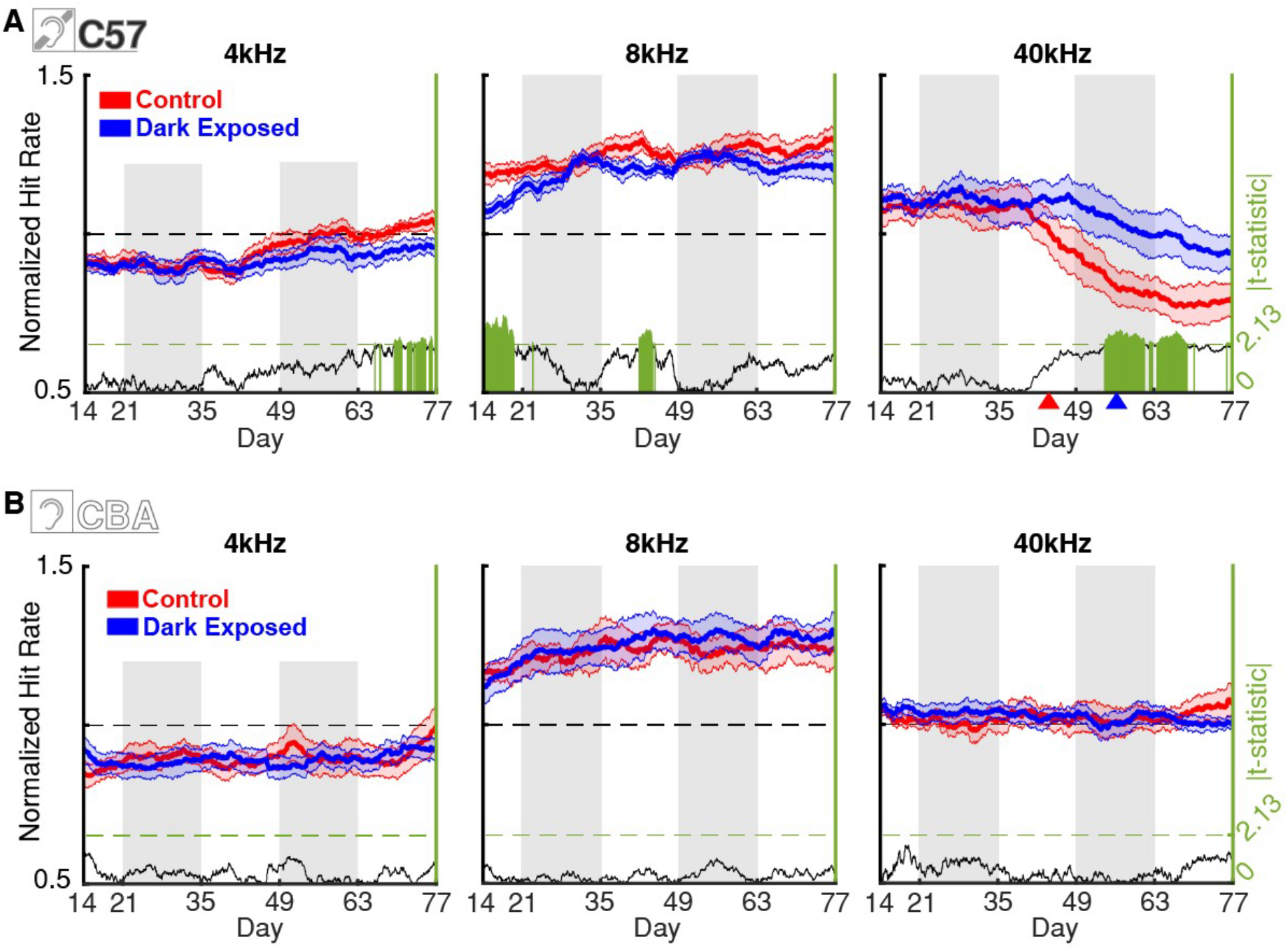
Rescuing effects of DE emerge in Period 3 of the C57BL/6J group. **(A)** (Left axis) Normalized hit rates for 4kHz, 8kHz, and 40kHz tones for CT C57BL/6J (red) and DE C57BL/6J (blue) ToneBoxes. Data averaged over all SPL levels. The shaded error bar represents the standard error of the mean. Red and blue arrowheads in the right panel for 40kHz mark the point in the timeline (days 45 and 57) where mean NHR deviated from the baseline by two standard deviations for CT and DE groups respectively. (Right axis) The absolute value of *t*-statistic from a two-sample t-test, if the *t*-test was performed at a given point in time. Green bars show data points of statistical significance (*t-test, p* < 0.05). **(B)** Same as in (A), but for the CBA CT and DE groups.

Together, these results suggest that DE increases performance on an auditory detection task in both CBA and C57BL/6J mice. However, the details in which DE benefits audition seem to differ between the two models: We hypothesize that in CBA mice with preserved hearing DE facilitates the improvement of tone detections across frequency ranges, while in C57BL/6J DE allows compensation to attenuate the age-dependent high-frequency hearing loss.

### DE delays the effects of presbycusis in C57BL/6J mice by 12 days

Our results suggest that C57BL/6J DE mice have higher behavioral performance at high sound frequencies compared to CT at the end of our experimental time window. We next aimed to identify the detailed behavioral time courses of these performance differences. We thus analyzed the changes in the sound-frequency-dependent performance in the CT and DE groups over time. For each ToneBox we calculated the normalized hit rate (NHR) for each stimulus (frequency & amplitude, FxA) condition in each hourly time bin for the entire 63 days of the experiment. We normalized the stimulus-specific (FxA) hit rates to the total hit rate of a given ToneBox for the same period across all conditions, meaning that the ToneBox with the normalized hit rate of 1 for a given FxA band had the same hit rate as all the FxA bands combined. This normalization enabled us to investigate the distribution of hit preference and ability across the sound spectrum. To minimize the effects of the 24-hour circadian rhythm, we averaged these normalized hit rates for both the CT and DE groups with a moving 168-hour window. NHR activity for C57BL/6J mice is shown for 4, 8, and 40kHz bands (Fig. 3A) and all remaining frequency bands (Fig. S2A). In C57BL/6J CT cages, a drop of the NHR for the 40kHz band is present starting around Day 40. In the C57BL/6J DE group, a much weaker drop is present at much older ages (Fig. 3A, right). To better define this difference in the onsets of performance decline for CT and DE groups, we defined the onset of decline as the first day when mean performance drops two standard deviations below baseline performance from period I. Respective days are labeled with colored arrows on the x-axis. This difference turned out to be 12 days (Day 45 vs 57). For the 4khz and 8kHz bands, no decreases in NHR were observed. Thus, DE delays the development of the behavioral effects of high-frequency hearing loss in C57BL/6J mice. Notably, significant differences in the 8kHz band were observed during period one around days 14-20. This was likely due to group fluctuations in the shaping phase and these differences ceased after day 20. Lastly, there was an increased preference for 4 and 6.3kHz bands in the CT group (Fig. S2) that occurred at the same time as the decline of the 40kHz band of this group. As discussed above, these increases were likely the compensation for lost ability in the 40kHz band.

In contrast to C57BL/6J mice, CBA mice do not suffer from any systemic peripheral hearing loss at this age. Consistent with this, our analysis shows that CBA CT and DE mice do not show differences in NHR in the 4, 8, and 40kHz bands (Fig. 3B) and all other frequencies (Fig. S2B). These results confirm that frequency preferences in CBA mice did not change with DE as was the case in the C57BL/6J group. Thus, the increase in absolute hit rates we observe in CBA mice (Fig. 2C, D) is relatively widespread across the frequency spectrum.

Together, these results support our hypothesis that DE selectively increases the relative performance of C57BL/6J mice at high frequencies while providing a more general benefit across a wide range of frequency spectrum in CBA mice.

### The effect of DE on performance is present across sound amplitudes

So far, we have lumped the performance at all sound amplitudes. We next evaluated if the increased performance of C57BL/6J DE mice was present at all sound amplitudes. We thus computed performance audiograms. We averaged cage NHRs for each period and plotted smoothed heatmaps for Period I and V of the C57BL/6J CT and DE groups (Fig. 4A; see Fig. S3 for data from all periods). The interquartile range (IQR) contours for high-frequency bands differ between C57BL/6J CT and C57BL/6J DE, and the CT group shows a shift of the IQR contours toward lower frequencies. To evaluate this further, we plotted the ratios of C57BL/6J DE vs. CT NHR (Fig. 4A bottom). The results from the C57BL/6J group indicate that DE effects are present across high-frequency bands (32 and 40kHz). A similar analysis from the CBA group shows very stable audiograms where the contours of the audiograms of the NHRs are nearly identical for Periods I and V (Fig 4B; see Fig. S4 for data from all periods). This is also visible in the ratios of CBA DE vs. CBA CT NHR (Fig. 4B bottom) where only minor differences are observed. We will next quantify these differences in detail.

**Figure 4:**
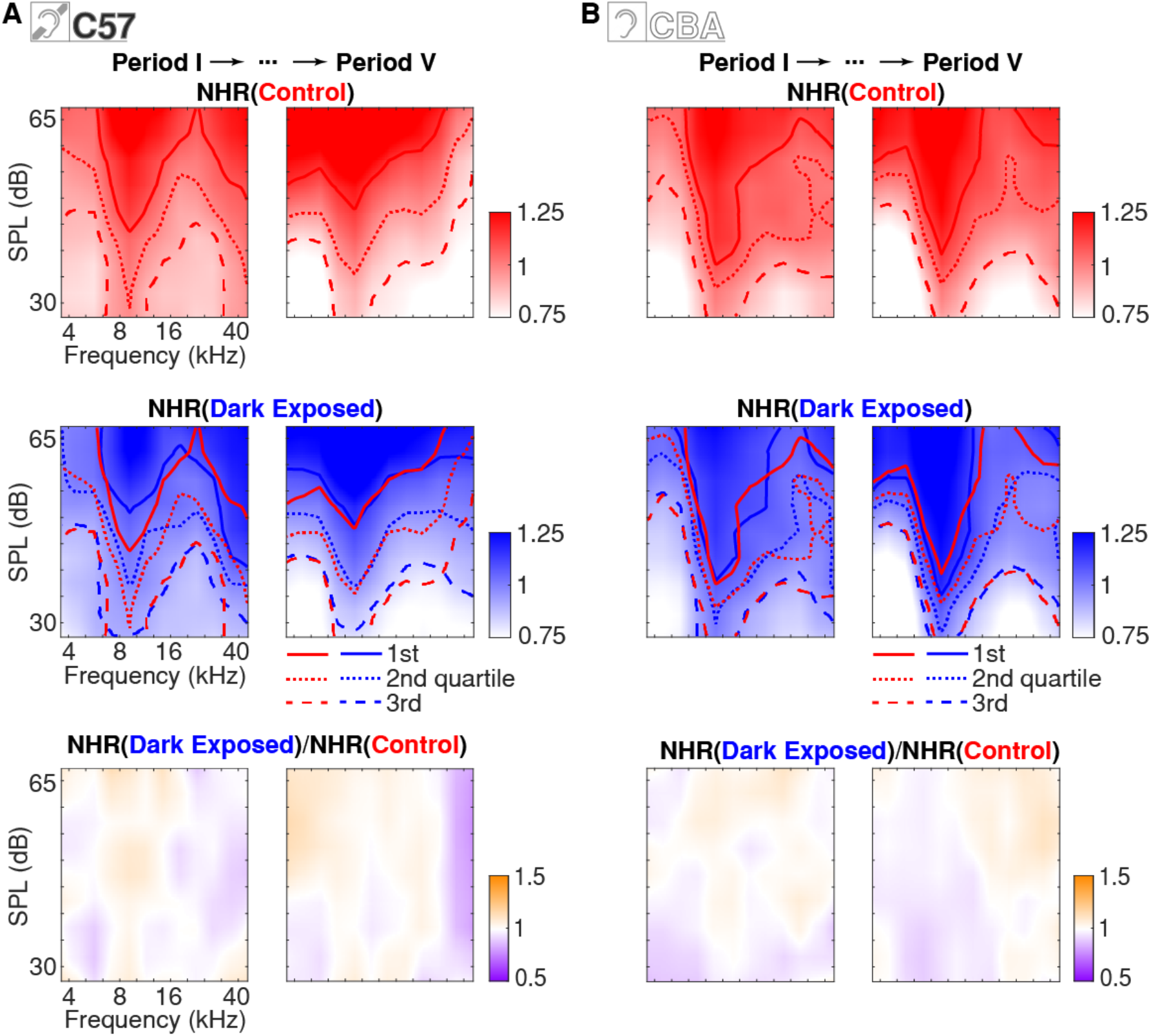
Frequency-amplitude-dependent performance is not altered by DE in CBA mice. **(A)** Normalized hit rate audiograms for all stimulus conditions for periods I (left) and V (right) of the C57BL/6J CT group (upper row) and C57BL/6J DE group (middle row). Dashed lines indicate quartiles of smoothed NHR. Red IQR lines for the DE group are plotted as CT reference. (lower row) The ratio of normalized hit rates between CT and DE C57BL/6J. **(B)** Same as in (A) but for the CBA CT and DE groups.

DE enhancement across performance levels suggested that DE had effects across tone amplitudes. We thus next investigated the DE enhancement effect in the amplitude domain by plotting the NHRs of the FxA bands along the amplitude dimension for two frequency bands of the C57BL/6J group from the period I and V: 8 and 40kHz, comparing the relationship between sound amplitude and NHR with our linear regression model as previously done on raw hit rate rations in Fig. 2E and F (Fig. 5A; see Fig. S5A for data from all periods and frequencies).

**Figure 5:**
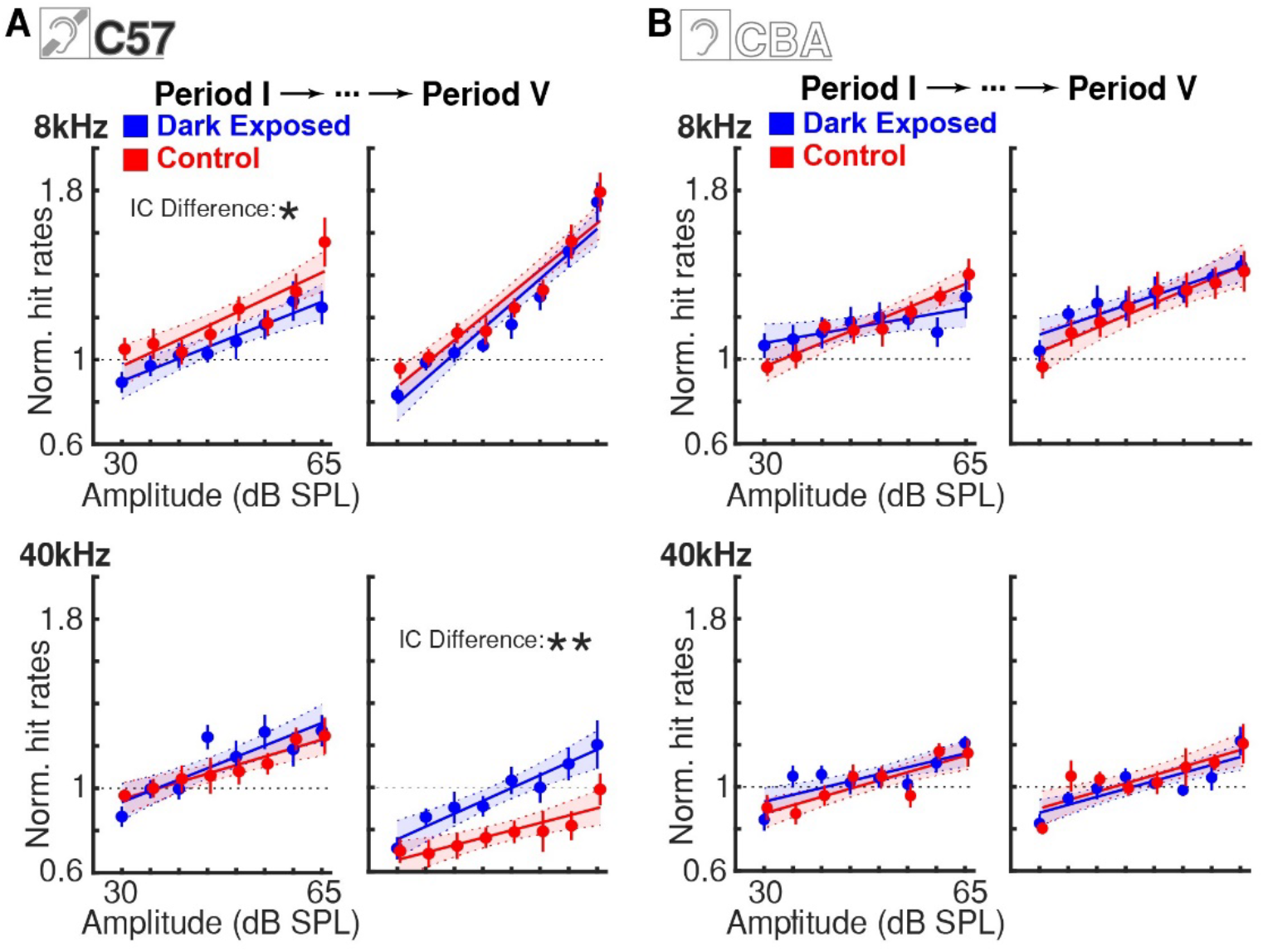
DE performance increases are multiplicative across levels. **(A)** Scatter points show normalized hit rates for 8kHz and 40kHz as a function of amplitude in the C57BL/6J CT (red) and DE (blue) group during Period I (left column) and Period V (right column). Vertical lines show SEM. Scatter points are overlapped with linear regression model fit in matching colors. Shaded areas are 95% confidence intervals of the fit. Intercept (IC) or slope (SL) Difference appears if the *p*-value of the *F*-test showed significance for either parameter of the group difference linear fit model (*F*-test, Bejnamini & Hochberg false discovery rate procedure applied. Does not appear for non-significant, * for *p* < 0.05, & ** for *p* < 0.01). Dashed black horizontal lines outline an NHR level of 1. **(B)** Same as in (A) but for CBA CT and DE groups.

As can be seen from Fig. S5A and Table S1, the significance of the difference between CT and DE groups for C57BL/6J mice is restricted to only 4 cases, 8kHz band in Period I that likely originated in group-wide fluctuations of performance in late days of the training (*F*-test, p=0.0465), 40kHz band for Period IV and V that signifies the rescuing effects of DE on 40kHz band (*F*-test, P.IV: *p* = 0.0003; P.V: *p* < 0.0001, and, lastly, 4kHz band in Period V which is compensatory effect of lost performance in the 40kHz band during the same period. For the last case, the slope parameter was also significant, meaning that CT mice increased their 4kHz band performance with increased gain compared to the DE group (F-test, IC: *p* = 0.0082; SL: *p* = 0.0117). In contrast, the CBA group revealed no significant differences between the two models (Fig. 5B and Fig. S5) for any frequency band or period (See Table S1).

Together, these analyses suggest that DE-induced auditory behavioral enhancement seen in the C57BL/6J group leads to decreased thresholds across the high-frequency spectrum after the onset of hearing loss.

### The effect of DE emerges gradually

We next investigated in detail the time course of the effects of DE. We thus generated NHR across the experimental periods for all stimulus combinations for both the C57BL/6J CT, C57BL/6J DE, CBA CT, and CBA DE groups and consequently plotted the ratios of CT vs. DE for both the C57BL/6J group (Fig. 6A) and CBA group (Fig. 6B). This analysis shows that the onset of DE enhancement for the C57BL/6J group emerged around day 40 for the softest high-frequency sounds and that improvements in the 32kHz bin emerged at around day 60. Given that day 40 was between our first and second DE periods, these data suggest that the first DE period could already have a long-lasting effect on tone detection performance.

**Figure 6:**
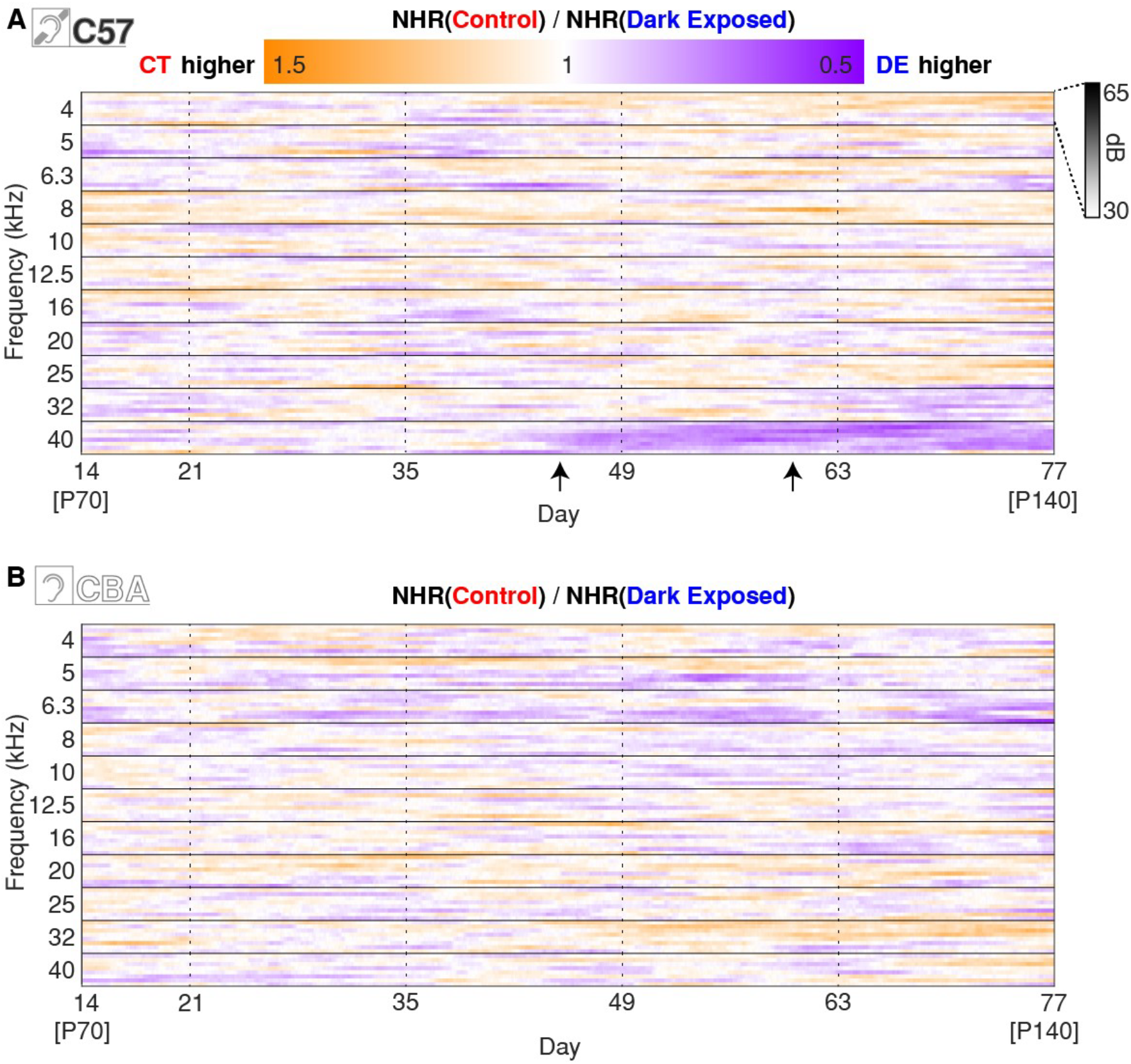
DE performance for quietest high-frequency tones increases after the first DE period. **(A)** The ratio of hit rates between CT and DE for all stimulus combinations of the C57BL/6J group. Arrows: Differences for 40kHz emerge after day 40, while differences for 32kHz emerge around day 65. Dashed vertical lines mark the individual periods. **(B)** Same as in (A) but for the CBA group.

To test if a single period of DE could have a preventative effect, we trained an additional cohort of C57BL/6J animals starting at P84 up to P140 (4 cages). Then, we performed two weeks of DE timed to match Period IV in a postnatal reference. A single period of DE also resulted in preventing the decreased performance in the 40kHz band (Fig. S6). Thus, a single period of DE was able to reduce the effect of presbycusis.

## Discussion

Our results show that temporary visual deprivation via DE in adults enhances the behavioral performance of C57BL/6J mice in tone detection tasks in high-frequency bands where the effects of presbycusis are usually evident. Additionally, we observed broad increases in the performance of low and mid frequencies in CBA mice that do not suffer from any systemic hearing loss at this age.

Our automated design allowed us to gather hundreds of thousands of trials per cage. Because of the minimalistic impact of the experiment design on mice’s daily routine, we eliminated several confounds commonly appearing in rodent behavior studies, such as repeated handling of animals (26). The hearing of both C57BL/6J and CBA mouse lines was previously studied extensively by several methods, most notably auditory brainstem response (ABRs) (19, 20, 22). Threshold intensity shifts caused by presbycusis in C57BL/6J mice were observed as early as P30 (27). While we did not measure the ABRs of individual mice, our experimental design minimized variability. First, we used large cohorts of animals enabled by our automatic system. Second, all animals were subject to the same developmental conditions until they were distributed to two experimental groups at the same age (P63) before the start of the experiment. Third, animals were group-housed in the ToneBox, thus each ToneBox recording represents a composite of the performances of the three individual mice within a given cage. This within-cage averaging further reduced the effect of the population variability on the hearing capabilities of individual animals.

While we here use a tone-detection task, the circuit and functional changes of DE are widespread and include the sharpening of tuning curves (10). We predict that DE affects a variety of auditory tasks. Indeed, training on auditory temporal discrimination tasks can also improve spectral tuning (28), suggesting that mechanisms engaged by training are affecting general sound processing. Given that DE has an effect on thalamic (14), thalamocortical (10), and intracortical (11-13) auditory circuits we expect that performance in a variety of auditory tasks is improved. Our data show an enhancement of the performance at high frequencies in C57BL/6J mice. This enhancement is consistent with the increased number of neurons responsive to high frequencies after DE in C57BL/6J mice (15).

C57BL/6J mice have early onset of presbycusis – gradual age-related hearing loss, evident in ABRs, otoacoustic emissions, and startle behavior for the high-frequency range starting around 10 weeks of age (18-20). When mice reached an age corresponding to when high-frequency ABR hearing threshold shifts were evident in this strain (19), we observed a decline in performance in our operand conditioning task in the high-frequency band. We noted that Control mice show an increase in the relative amount of hits to low frequencies, indicating that they shift their behavior to relatively “easier” stimuli to keep their water consumption constant. In contrast, DE mice showed better performance at high frequencies. What mechanism could underlie this improved performance? Hearing loss in C57BL/6J is caused by degeneration of the cochlea and this degeneration starts at high frequencies (22-24). Such degeneration results in reduced ascending sound-evoked activity and subsequently reduced activation of the auditory cortex for high-frequency stimuli (29). Mechanistically, our behavioral data could be explained by the circuit level plasticity we reported in previous studies. DE induces an increase in the strength of auditory thalamocortical synapses (10), which can counteract the reduced afferent drive which leads to enhanced sensitivity and increased responsiveness to sound stimuli in the thalamocortical recipient layers of the auditory cortex (10). Consistent with the increase in thalamocortical synaptic gain, we observed that DE leads to a steep increase in firing rates with changes in sound amplitudes (10). Intracortical circuits and thalamic circuits can alter gain and adult DE has been shown to affect both. After DE, ascending and recurrent intracortical circuits change synaptic strength (11-13), and refine their connections, leading to a more efficient information transmission (12). In addition to changes on the cortical and thalamocortical level, DE reduces inhibition from the thalamic reticular nucleus to the auditory thalamus, enhancing the ascending transmission of sound information through the thalamus (14). DE could also have effects on spectral contrast tuning sensitivity (30). As previously shown, DE induces decorrelation of the sound-evoked population activity in A1 which can lead to increased encoding fidelity of represented stimuli in the cortex (15, 31). Together, all these circuit changes, both at the thalamic and cortical levels, enhance the transmission of the weakened high-frequency ascending signals to the auditory cortex and lead to an increased representation of high-frequency tones in the auditory cortex after DE (15). This potentiation of the feedforward circuit and refinement of the intracortical circuit could allow better detection and sharper tuning needed to allow the processing of reduced auditory signals arising from age-related peripheral hearing loss. We reason that these extensive circuit changes compensate for decreased ascending drive and lead to the observed reduced behavioral performance declines after DE.

Studies investigating early and life-long visual deprivations have shown various functional and circuit changes that can give rise to improved auditory performance (32-34). We find that DE can also induce such changes in adult animals. The changes we here see with DE improve auditory behavior in a model of presbycusis is consistent with the idea that the behavioral deficits in presbycusis are not solely due to loss of inner hair cells in the cochlea but also due to changes in the brain. Indeed, the aging auditory cortex in CBA mice that do not suffer from peripheral hearing loss also shows altered sound-evoked activity, such as increased correlations and reduced ability to control activity correlations (35, 36). Our observation of increased hit rates in DE CBA mice across low- and mid-frequencies is consistent with these findings. In conclusion, this study suggests that the changes in central auditory processing lead to the increased ability of animals to perform auditory tasks after DE. Furthermore, our data collectively suggest that DE could be a simple method to reduce some of the effects of central aging and enhance the efficacy of auditory performance with cochlear implants.

## Acknowledgments

We thank members of the Kanold lab for their comments on the manuscript.

We thank Dr. Behtash Babadi for his advice regarding interpreting the results of the manuscript.

## Author contributions

Conceptualization: POK, HKL, PJ

Methodology: PJ, POK

Investigation: PJ

Visualization: PJ, POK

Funding acquisition: POK, HKL

Project administration: PJ, POK

Supervision: POK

Writing – original draft: PJ, POK

Writing – review & editing: PJ, HKL, POK

## Data and materials availability

All data supporting the findings from this study will be available upon publication at the Johns Hopkins Data Archive (https://archive.data.jhu.edu).

## Supplementary Materials

Materials and Methods

Supplementary Figures S1-S6

Supplementary Table S1

