## Supplementary Materials for "Dark exposure reduces high-frequency hearing loss in C57BL/6J mice"

**Materials and Methods**

**Supplementary Figures S1-S6**

**Supplementary Table S1**

#### Materials and Methods

All experimental procedures were approved by the Johns Hopkins University Animal Care and Use Committee.

##### Animals

60 (30m/30f) C57Bl/6J mice (Jackson Laboratory strain #000664) and 48 (24m/24f) CBA/CaJ mice (Jackson Laboratory strain #000654) were raised in a 12h light /12h dark cycle. At the beginning of postnatal week 10, mice were randomly allocated to the behavioral chambers for the initial training. All animals survived for the duration of the whole experiment. Food was provided ad libitum. The weight of the animals remained nominal throughout the duration of the experiment, similar to our previous studies (1, 2). The bedding was changed before every period during the experiment (once every 14 days). Mice were checked for any injuries or effects of water restriction, such as skin turgor with no reports for the duration of the experiment.

##### ToneBox & sound calibration

We based our home-caged based behavior station on our previously published ToneBox design (1, 2). The ToneBox is a device composed of a behavioral interface (BI) comprised of a 3D printed housing containing a speaker (PUI Audio, Inc., AS02708CO-WR-R), its amplifier (Adafruit, Mono 2.5W Class D Audio Amplifier, PAM8302), 3d printed lick spout, and the wiring cables for the delivery of the DC power supply, speaker analog signal, and the capacitance signal which is connected to the licking spout. This device is connected to the central control unit (CCU) outside of the enclosure box, composed of a Raspberry Pi computer (Raspberry Pi 3 Model B+), a custom-made GPIO PCB interface, including a capacitance sensor (Adafruit, AT42QT1010), and a USB sound card (Steinberg, UR12, 192kHz sampling rate). The system operation is controlled by a Python script executed on the Raspberry PI. 8 ToneBoxes are located on a ToneBox rack, and the cluster of 8 Raspberry PIs is connected to a monitoring PC via an Ethernet switch. Once every 24 hours, data is transferred to the monitoring PC to evaluate the animals' performance.

Mice receive a water reward after successfully activating the capacitance sensor within 3 seconds of the tone onset. This is the only source of water, except for a burst of free water (equivalent to 50 hit trials) occurring every 17-19 hours. This free reward is intentionally desynchronized from the Circadian cycle and secures minimal survival water intake. The water burst also prevents clogging of the water lines and ensures reliable long-term water delivery.

The BI, CCU, and home cage were individually placed within a sound-attenuating chamber composed of a cabinet with a clear door and a ventilation fan. The chambers provide >20 dB attenuation between ToneBoxes for tones >1 kHz, allowing simultaneous auditory training.

Tones were generated and calibrated for each ToneBox using a custom-written Python script. There were 11 frequency levels between 4 and 40kHz (1/3$^{rd}$ octave spacing: 4, 5, 6.3, 8, 10, 12.5, 16, 20, 25, 32, 40 kHz) and eight amplitude levels between 65 and 30dB SPL (5dB spacing), resulting in 88 different sound stimuli randomly presented through the speaker. Tones were calibrated using a pre-polarized ¼ inch microphone (Brüel and Kjær, TYPE 4944) and signal preamplifier (Brüel and Kjær, TYPE 1704-A-002). During calibration, a microphone was placed directly under the main axis of the ToneBox speaker, approximately at the height where mice heads would be located if standing under the device. A signal from the preamplifier was routed to the UR12 sound card and further processed in Raspberry Pi by a custom-written Python script. 65dB tones of each frequency were calibrated with a precision of ± 1dB SPL compared to the 1kHz 94dB sound calibrator. The rest of the amplitude levels were attenuated by 5dB levels. A bandpass filter was applied to the recorded waveform before calculating sound RMS (root mean square) amplitude. The bandpass filter had high-pass and low-pass frequencies of 0.95 and 1.05 times the frequency of the tone currently calibrated.

The base Inter-trial interval was randomized between 17 and 21 seconds. However, if mice licked the water spout before the onset of the sound, the inter-trial timer reset to 0. This setting forces mice from random licking towards tone-detection behavior. Our behavioral system does not detect the performance of individual mice. Events from the Tonebox devices were saved as time-stamped events (30ms resolution, ~33Hz sampling rate) locally to the flash drive of Raspberry PI. Sound ID, trial result, trial inter-trial intervals, and capacitance licking data were recorded for the duration of the experiment.

Next, we will clarify different values of raw hit rates in Fig. 1B, 1E-H, and Fig. S1. Figures 1B and S1 reported hit rates of ~20-40/hour because they report a full performance, while figures 1E-1H show frequency-dependent performance, thus on average these values are 1/11 of the full performance since 11 different frequencies are being tested.

Visual deprivation settings

On the day of switching from the 12h Light / 12h Dark setting to the 24h Dark Exposure setting, chambers were sealed from any light source by a triple barrier of blackout curtains made of high-density fabric material. First, chamber windows were sealed with the initial layer; then, a second layer was placed over the entire row of 4 chambers in each rack, followed by a final layer covering both rows combined and finally covering up the whole rack. Mice's licking behavior was monitored remotely, and no anomalies were detected in our experiments.

Data analysis

For the "continuous" timeline figures, we calculated histograms with 1h bins of hit counts ($hc_{fa}$) and trial counts ($tc_{fa}$) for each frequency x amplitude tone stimulus. We calculate the raw hit rate $rhr$ as

$$rhr_{fa} = \frac{hc_{fa}}{tc_{fa}}$$

for a given time bin. Additionally, for the same 1h time bins, we calculate total hit counts ($thc$) and total trial counts ($ttc$), irrespective of the sound stimulus. We then calculate the normalization factor ($nf$) for each bin as

$$nf = \frac{thc}{ttc}$$

The normalized hit rate ($nhr_{fa}$) is then calculated as:

$$nhr_{fa} = \frac{rhr_{fa}}{nf}$$

$nhr_{fa} = 1$ means that the cage performs at a given frequency x amplitude on average same as the time interval total hit rate. Values $> 1$ show increased frequency x amplitude performance, while values $< 1$ show performance that is lower than the total performance. To remove oscillations caused by the circadian rhythm, a moving average with a window of 168h (7 days) was applied. A similar computation was performed for the calculation of audiograms, except that time bins were increased to the duration of the whole period, which was 336h (14 days).

For testing of statistical significance, we used a two-sample, two-tailed *t*-test, *a*=0.05, *p*>=0.05, *p*<0.05, *p*<0.01 labeled as "n.s.", "*", and "**" respectively.

#### Supplementary Figures

## (S1 – S6)

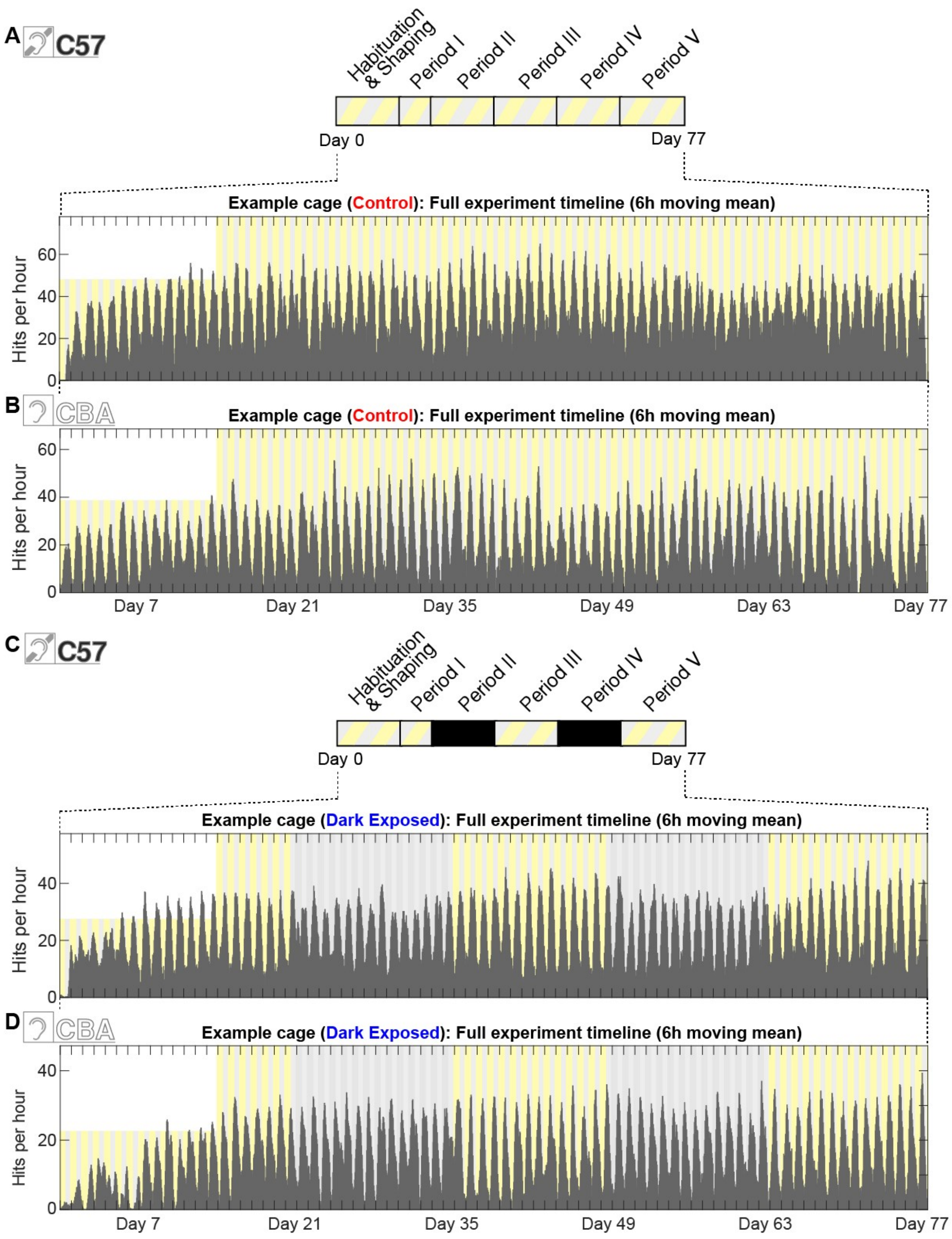
A
C57
Habituation & Shaping
Period I
Period II
Period III
Period IV
Period V
Day 0
Day 77
Example cage (Control): Full experiment timeline (6h moving mean)
Hits per hour
B
CBA
Example cage (Control): Full experiment timeline (6h moving mean)
Hits per hour
Day 7
Day 21
Day 35
Day 49
Day 63
Day 77
C
C57
Habituation & Shaping
Period I
Period II
Period III
Period IV
Period V
Day 0
Day 77
Example cage (Dark Exposed): Full experiment timeline (6h moving mean)
Hits per hour
D
CBA
Example cage (Dark Exposed): Full experiment timeline (6h moving mean)
Hits per hour
Day 7
Day 21
Day 35
Day 49
Day 63
Day 77

**Supplementary Figure 1: Cage examples of hit rates for the entire experimental timeline.**

**(A)** Example timeline of hourly hit rates for the C57BL/6J Control cage, **(B)** CBA Control cage, **(C)** C57BL/6J DE cage, and **(D)** CBA DE cage for the duration of the whole experiment, including habituation and shaping phase. 6-hour moving average applied. Day 0 corresponds to the postnatal day (P 63). Black shading in Period II and IV and between days 21-35 and 49-63 for the DE show when the Dark Exposure took place.

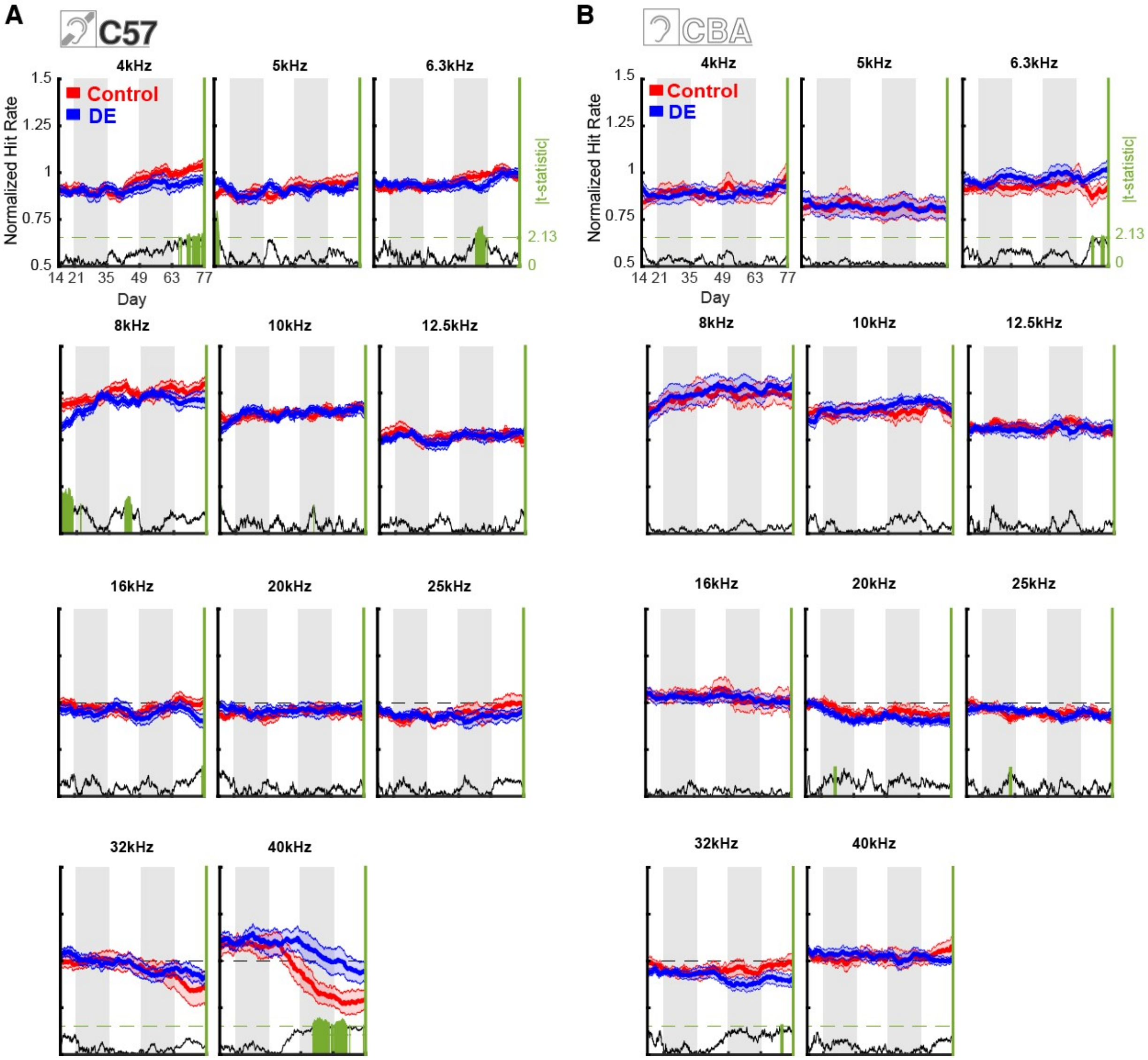


**Supplementary Figure 2: Timeline comparison of NHR for all frequency bands of C57BL/6J and CBA groups.**

**(A)** Timeline of the normalized hit rate (NHR) averaged across the amplitudes for all frequency bands. (Left axis) Normalized hit rates for C57BL/6J CT (red) and DE (blue) ToneBoxes. Data averaged over all SPL levels. Shaded areas show SEM. (Right axis) The absolute value of t-statistic from two-sample t-test. Green bars show data points of statistical significance (t-test, $p<0.05$). **(B)** Same as in (A), but for the CBA group.

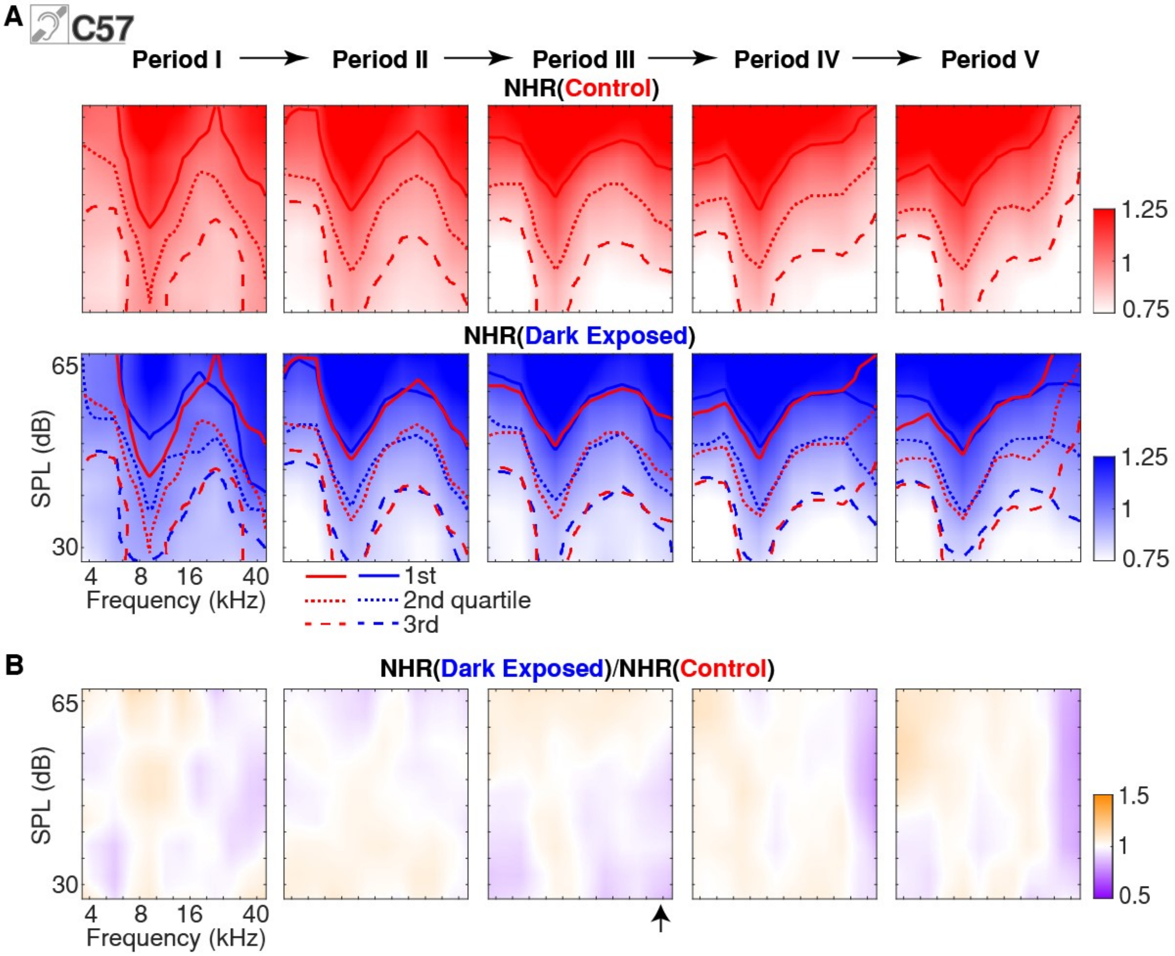


**Supplementary Figure 3: Rate-of-change audiograms for all periods of C57BL/6J group.**

**(A)** Normalized hit rate audiograms for all stimulus conditions and all periods of the C57BL/6J CT group (upper row) and C57BL/6J DE group (lower row). Dashed lines indicate quartiles of smoothed NHR. Red IQR lines for the DE group are plotted as CT reference. **(B)** The ratio of normalized hit rates between CT and DE C57BL/6J.

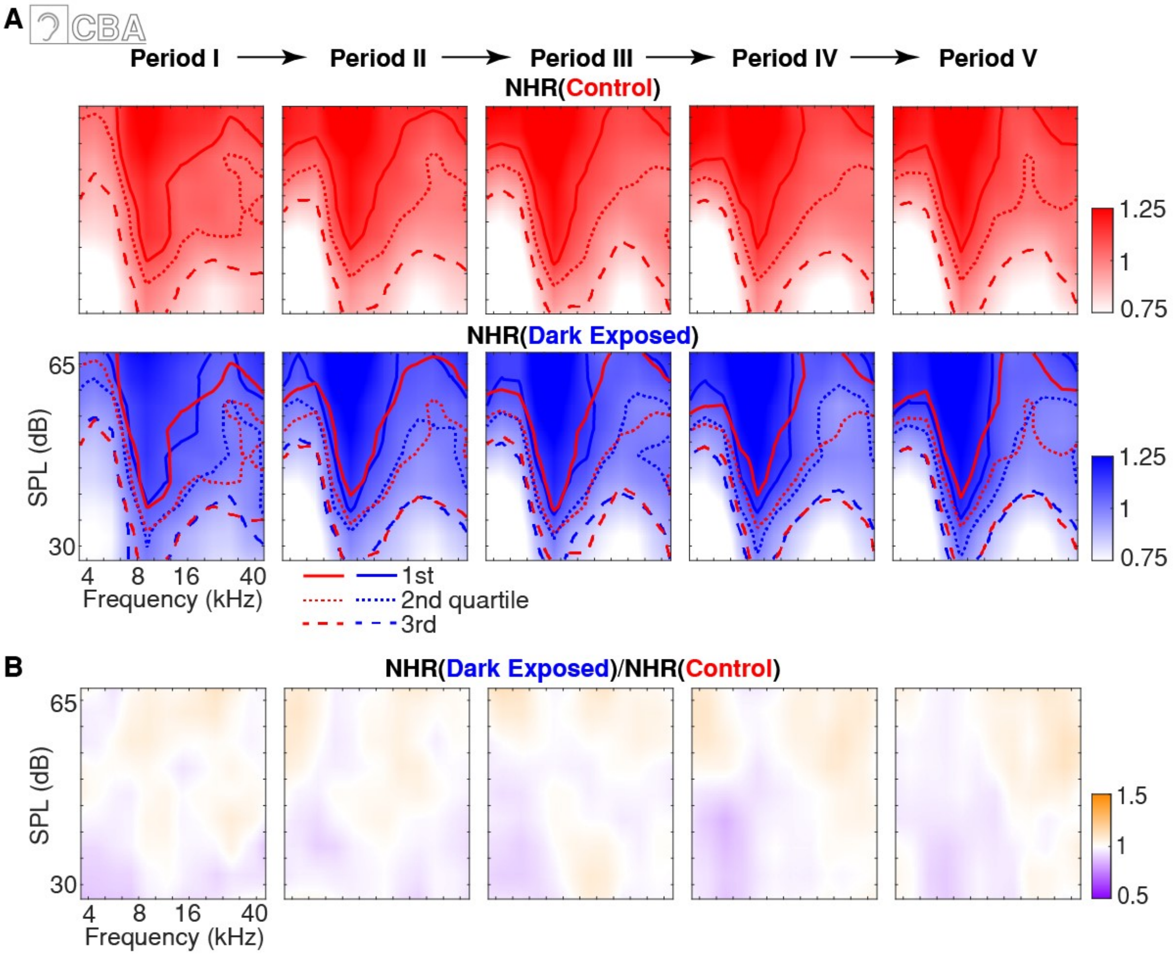


**Supplementary Figure 4: Rate-of-change audiograms for all periods of the CBA group.**

**(A)** Normalized hit rate audiograms for all stimulus conditions and all periods of the CBA CT group (upper row) and CBA DE group (lower row). Dashed lines indicate quartiles of smoothed NHR. Red IQR lines for the DE group are plotted as CT reference. **(B)** The ratio of normalized hit rates between CT and DE CBA.

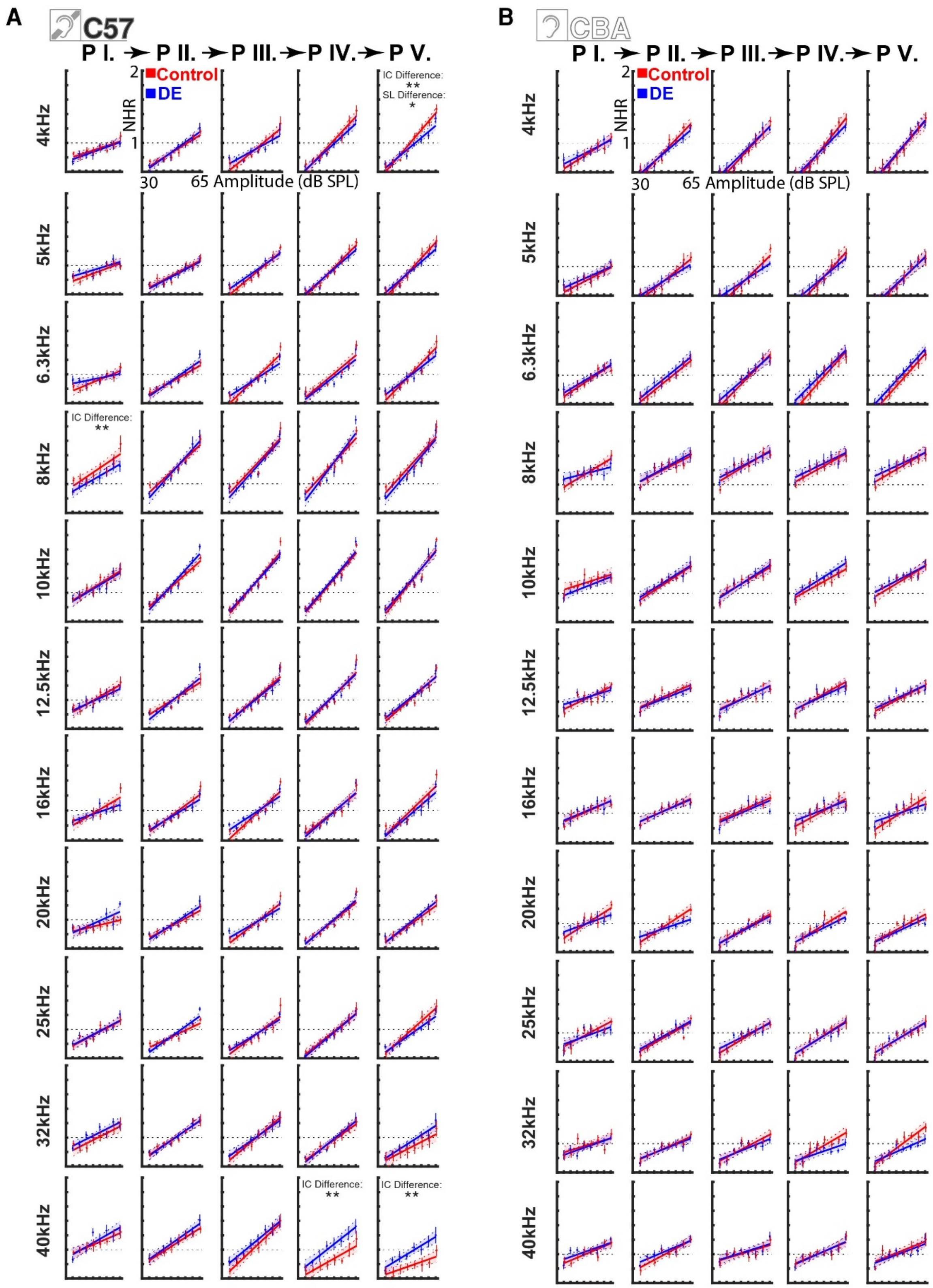
A
C57
P I. ➤ P II. ➤ P III. ➤ P IV. ➤ P V.
Control
DE
NHR
2
1
30
65 Amplitude (dB SPL)
IC Difference: **
SL Difference: *
IC Difference: **
4kHz
5kHz
6.3kHz
8kHz
10kHz
12.5kHz
16kHz
20kHz
25kHz
32kHz
40kHz
B
CBA

**Supplementary Figure 5: Amplitude-dependent NHRs for all frequency bands of the C57BL/6J and CBA groups.**

**(A)** Scatter points show normalized hit rates for all tested frequencies as a function of amplitude in the C57BL/6J CT (red) and DE (blue) group during Period I (most left column) to Period V (most right column). Vertical lines show SEM. Scatter points are overlapped with linear regression model fit in matching colors. Shaded areas are 95% confidence intervals of the fit. Intercept (IC) or slope (SL) Difference appears if the *p*-value of the *F*-test showed significance for either parameter of the group difference linear fit model (*F*-test, Bejnamini & Hochberg false discovery rate procedure applied. Does not appear for non-significant, * for $p < 0.05$, & ** for $p < 0.01$). Dashed black horizontal lines outline an NHR level of 1. **(B)** Same as in (A) but for the CBA group. The overview of *p*-values is provided in Supplementary Table 1.

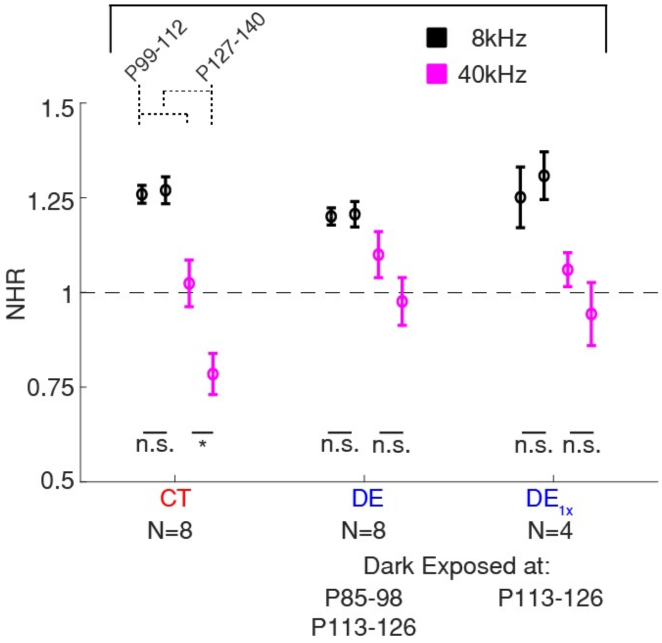


**Supplementary Figure 6: A single period of DE can rescue high-frequency performance.**

A single session of DE was performed on 12 C57BL/6J mice (4 cages, 6m, 6f), timed depending on the postnatal age of the animals. Four cages were exposed to a single period of DE during P113-126 (corresponding to Period IV) ($DE_{1x}$). Plotted are averaged NHR for 8kHz (black) or 40kHz (magenta) in the two-week-long periods. Error bars show SEM. 40kHz NHR of $DE_{1x}$ was similar to the NHR after 2 DE periods. 40kHz CT band was the only one that showed a significant difference between the two observed periods. ‘*’ indicates statistical significance, n.s. not significant (t-test, $p<0.05$).

### Supplementary Table

# (S1)

| | | P1 | P2 | P3 | P4 | P5 | | P5/P1 |
|---|---|---|---|---|---|---|---|---|
| | C57BL/6J Intercept (*p*-value adjusted) | | | | | | | C57BL/6J Intercept (*pva*) |
| Frequency (kHz) | 4.0 | 0.6211 | 0.9188 | 0.9188 | 0.2197 | 0.0082 | | 0.9552 |
| | 5.0 | 0.3720 | 0.6211 | 0.6000 | 0.6211 | 0.6000 | | 0.9552 |
| | 6.3 | 0.6211 | 0.6487 | 0.6487 | 0.2744 | 0.6211 | | 0.9725 |
| | 8.0 | 0.0465 | 0.6487 | 0.2825 | 0.6211 | 0.0963 | | 0.6843 |
| | 10.0 | 0.6487 | 0.6211 | 0.6487 | 0.6487 | 0.9188 | | 0.8520 |
| | 12.5 | 0.6487 | 0.8545 | 0.6000 | 0.9188 | 0.7688 | | 0.8520 |
| | 16.0 | 0.6281 | 0.6000 | 0.6000 | 0.6211 | 0.0651 | | 0.9552 |
| | 20.0 | 0.4205 | 0.6487 | 0.9932 | 0.9188 | 0.6211 | | 0.9552 |
| | 25.0 | 0.9188 | 0.6487 | 0.9834 | 0.6211 | 0.0963 | | 0.9552 |
| | 32.0 | 0.2825 | 0.7699 | 0.9564 | 0.9564 | 0.2825 | | 0.9552 |
| | 40.0 | 0.6487 | 0.6000 | 0.2825 | 0.0003 | 0.0000 | | 0.0027 |
| | C57BL/6J Slope (*p*-value adjusted) | | | | | | | C57BL/6J Slope (*pva*) |
| Frequency (kHz) | 4.0 | 0.8818 | 0.5938 | 0.1134 | 0.3639 | 0.0117 | | 0.9431 |
| | 5.0 | 0.6952 | 0.9746 | 0.3351 | 0.3639 | 0.1316 | | 0.9431 |
| | 6.3 | 0.2714 | 0.3639 | 0.0182 | 0.7907 | 0.1134 | | 0.9431 |
| | 8.0 | 0.6952 | 0.3410 | 0.8275 | 0.1134 | 0.6581 | | 0.9431 |
| | 10.0 | 0.9349 | 0.1134 | 0.5397 | 0.9746 | 0.6226 | | 0.9772 |
| | 12.5 | 0.6226 | 0.2714 | 0.7784 | 0.6226 | 0.6226 | | 0.9772 |
| | 16.0 | 0.2714 | 0.4398 | 0.0182 | 0.6226 | 0.5397 | | 0.9431 |
| | 20.0 | 0.3639 | 0.4104 | 0.1526 | 0.6311 | 0.6226 | | 0.9431 |
| | 25.0 | 0.9746 | 0.0182 | 0.4319 | 0.8818 | 0.2714 | | 0.9431 |
| | 32.0 | 0.9746 | 0.8818 | 0.4763 | 0.6226 | 0.6226 | | 0.9431 |
| | 40.0 | 0.6226 | 0.7534 | 0.6226 | 0.4319 | 0.2503 | | 0.9431 |
| | CBA Intercept (*p*-value adjusted) | | | | | | | CBA Intercept (*pva*) |
| Frequency (kHz) | 4.0 | 0.8361 | 0.8163 | 0.8361 | 0.8361 | 0.9692 | | 0.0115 |
| | 5.0 | 0.8163 | 0.9278 | 0.9692 | 0.9692 | 0.9815 | | 0.1292 |
| | 6.3 | 0.8361 | 0.8163 | 0.8361 | 0.6460 | 0.4476 | | 0.0014 |
| | 8.0 | 0.9278 | 0.9278 | 0.9278 | 0.8278 | 0.8278 | | 0.0079 |
| | 10.0 | 0.5695 | 0.8361 | 0.9815 | 0.5695 | 0.8278 | | 0.0005 |
| | 12.5 | 0.9278 | 0.8163 | 0.9278 | 0.8278 | 0.8361 | | 0.0107 |
| | 16.0 | 0.9278 | 0.9278 | 0.8163 | 0.8361 | 0.9815 | | 0.0071 |
| | 20.0 | 0.9278 | 0.6460 | 0.8278 | 0.6725 | 0.8163 | | 0.0807 |
| | 25.0 | 0.8163 | 0.8361 | 0.8361 | 0.9013 | 0.9815 | | 0.0107 |
| | 32.0 | 0.8361 | 0.8361 | 0.8163 | 0.4476 | 0.2409 | | 0.2082 |
| | 40.0 | 0.8163 | 0.8163 | 0.9278 | 0.9387 | 0.8163 | | 0.1292 |
| | CBA Slope (*p*-value adjusted) | | | | | | | CBA Slope (*pva*) |
| Frequency (kHz) | 4.0 | 0.8745 | 0.6408 | 0.9621 | 0.6408 | 0.9621 | | 0.7628 |
| | 5.0 | 0.9621 | 0.6408 | 0.2816 | 0.6408 | 0.9621 | | 0.9039 |
| | 6.3 | 0.9621 | 0.9824 | 0.8745 | 0.8745 | 0.9621 | | 0.9151 |
| | 8.0 | 0.2010 | 0.9621 | 0.9621 | 0.9621 | 0.9621 | | 0.6479 |
| | 10.0 | 0.9621 | 0.9621 | 0.9621 | 0.9621 | 0.9621 | | 0.6479 |
| | 12.5 | 0.6536 | 0.9621 | 0.9621 | 0.9621 | 0.9621 | | 0.7628 |
| | 16.0 | 0.9621 | 0.9621 | 0.9621 | 0.8745 | 0.2010 | | 0.6479 |
| | 20.0 | 0.4440 | 0.0713 | 0.9621 | 0.8745 | 0.9621 | | 0.6479 |
| | 25.0 | 0.6408 | 0.9621 | 0.8745 | 0.9621 | 0.9621 | | 0.6479 |
| | 32.0 | 0.9621 | 0.9621 | 0.8745 | 0.6151 | 0.1570 | | 0.6479 |
| | 40.0 | 0.9621 | 0.8745 | 0.9621 | 0.9621 | 0.9621 | | 0.8328 |

**Supplementary Table 1: FDR-corrected *p*-values overview for Figure 5 and Supplementary Figure S5.**

The table shows FDR-corrected p-values of the F-tests of the linear regression model fit estimating Intersect and Slope parameters. C57BL/6J Intercept and Slope panels are for Fig. 5A and Fig S5A, while CBA Intersect and Slope panels are for Fig. 5B and Fig. S5B. Right column containing C57BL/6J Intersect and Slope and CBA Intersect and Slope panes are for Fig. 5C and 5D. Dark and bright green background colorings highlight p-values that are *- and **- significant respectively (*F*-test, Bejnamini & Hochberg false discovery rate procedure applied.) The *pva* abbreviation in the narrow column stands for “*p*-value adjusted”.
